# Technical upgrade of an open-source liquid handler to support bacterial colony screening

**DOI:** 10.1101/2023.02.28.530165

**Authors:** Irene del Olmo, Pablo Yubero, Álvaro Gómez-Luengo, Juan Nogales, David R. Espeso

## Abstract

The optimization of genetically engineered biological constructs is a key step to deliver high-impact biotechnological applications. The use of high-throughput DNA assembly methods allows the construction of enough genotypic variants to successfully cover the target design space. This, however, entails extra workload for researchers during the screening stage of candidate variants. Despite the existence of commercial colony pickers, their high price excludes small research laboratories and budget-adjusted institutions from accessing such extensive screening capability. In this work we present COPICK, a technical solution to automatize colony picking in an open-source liquid handler Opentrons OT-2. COPICK relies on a mounted camera to capture images of regular Petri dishes and detect microbial colonies for automated screening. COPICK’s software can then automatically select the best colonies according to different criteria (size, color and fluorescence) and execute a protocol to pick them for further analysis. Benchmark tests performed for *E. coli* and *P. putida* colonies delivers a raw picking performance over pickable colonies of 82% with an accuracy of 73.4% at an estimated rate of 240 colonies/h. These results validate the utility of COPICK, and highlight the importance of ongoing technical improvements in open-source laboratory equipment to support smaller research teams.

## 1 Introduction

Recent developments in genome editing tools and protocols (i.e., Golden Gate (Engler et al., 2008), MoClo (Weber et al., 2011) or BASIC (Storch et al., 2015; Storch et al., 2017)) have enabled molecular biologists to build modular DNA sequences with an unprecedented velocity and success rate (McGuire et al., 2020). As a consequence, the assembly of complex functional genetic devices, such as artificial operons (Hérisson et al., 2022) or logic circuits (Jones et al., 2022) has become technically and economically accessible (Storch et al., 2020). The inherent tendency of such constructs to exhibit unpredictable outcomes due to their intrinsic biological complexity (Gardner, 2013) can be compensated by the high-throughput assembly capability that these techniques may deliver. However, the effort required to test all candidate constructions grows as well (Leavell et al., 2020). Typical clone screening stage requires the separation of cells by plating onto a solid substrate, followed by a further discrimination step between negative and positive colonies. Then, a user chosen selection criteria is applied to select the ones preferred for final phenotypic validation and genomic sequencing analysis.

This process can be performed manually in those cases where displaying the target phenotype involves low metabolic burdening, and the number of DNA subunits required to assemble designed genetic sequence is reduced. On the contrary when genetic design grows in complexity (i.e. large number of subunits, large concatenation of preassembled modules, etc.) and functionality (regulation), the combinatorial possibilities required to cover the whole design parametric space skyrockets, and the screening workload scales up above allowable values for human labor (Appleton et al., 2017; Leavell et al., 2020). At this point, the automation of selection and screening becomes the unique viable solution in terms of time and resource consumption (Chao et al., 2017; Hillson et al., 2019).

Colony pickers are well developed technical alternatives available on the market that alleviate the mentioned bottleneck in colony screening (Moffat et al., 2021). These devices consist of a processing unit that synchronizes two hardware blocks: an imaging system that captures an image of the solid plate (where colonies grow), and a robotic platform that allows the physical interaction between a picking tip and the colony plate. Prior to picking, a specialized software processes the image to detect, locate and characterize the colonies in the plate. This enables the discrimination and selection of the best colonies according to flexible criteria: colony size, geometry, color, luminescence or fluorescence (Stephens et al., 2002; Fabritius et al., 2018).

Despite the successes of these solutions, two major drawbacks have prevented them from being a common tool in microbiology laboratories. First, this technology was originally developed to cover the needs of high-edge research institutions and companies (Hansen, 1993). The design features, work capabilities and price of commercial units are fitted for customers dealing with heavy workloads. As a consequence, these devices are oversized (in terms of features and capacities) with respect to those required by smaller laboratories. Furthermore, the extra capabilities incur in an unaffordable price for such small customer profiles. Second, most commercial brands have developed their own programming ecosystems, enforcing customers to hire specialized technicians to operate the equipment, investing on costly training courses for their own staff or paying the manufacturer under demand for the design of custom-made protocols. Again, small actors are excluded from expanding their screening capabilities because of their lack of financial resources.

However, latest advances in open-source software and hardware may turn aside such situation (Villanueva-Cañas et al., 2021; Rodrigues et al., 2022; Sanders et al., 2022). The release of Opentrons, the first open-source liquid handler, has promoted the use of robotic automation in many laboratories due to its affordable price (around 7k$) and its accessibility via Python API (Villanueva-Cañas et al., 2021). Taking advantage of its native programming environment, a custom-made software package can be developed to couple hardware actuation in robot with analysis software specifically designed to infer colony positions. The use of convolutional neural networks (CNNs) has already demonstrated their potential to segment objects of interest within images provided that a good training stage is performed. A good example of this application is Detectron, an AI based software system designed by Facebook development team (Wu, 2019). It was successfully applied to different contexts, such as the profiling of Facebook accounts to create targeted propaganda (Rosenberg, 2018) or the analysis of medical images (Amerikanos and Maglogiannis, 2022). This inference model, released on GitHub under an open-source license in 2018, offers a good opportunity for community-driven applications to exploit its powerful features in other fields of study.

Here we present COPICK, a technical modification of the native OT-2 framework to integrate on-board image acquisition with a Detectron 2 based inference motor for online colony segmentation. Pixel-based inference is screened according to user criteria and mapped to physical domain, empowering OT-2 onboard pipette to pick individual colonies from solid-plate media. The content of this manuscript is presented as follows: the description of COPICK technical solution is detailed in section 2, which is subdivided in 8 parts. Subsection 2.1 presents the technical details of hardware adaptation. Subsections 2.2 to 2.4 describe the inference algorithm, the dataset preparation and the training and optimization process of the model. Subsections 2.5 to 2.7 covers the integration of the solution with OT-2 robot: colony screening methods, spatial allocation of detected segments and orchestrator software. Subsection 2.8 describe the benchmark tests performed to validate the platform. Results of validation tests are discussed in section 3. Section 4 presents an extended discussion about the technical tradeoffs and limitations of COPICK, together with some possible solutions to improve its performance. Finally, we retrieve some conclusions in section 5.

## 2 Method

### 2.1 Hardware adaptation

COPICK relies on the following hardware components:

1. A camera, to capture the spatial distribution of colonies.
2. A lighting gadget to improve the quality of captured snapshots.
3. A CPU that processes the image, segments and chooses a defined number of colonies according to several features and outputs their corresponding coordinates.
4. A computer numerical control (CNC) hardware in charge of physically accessing colonies to selectively pick biomass and transfer it to a target destination.

This technical upgrade was designed to be implemented into an Opentrons OT-2, an automated liquid-handler platform that can execute custom protocols with a predefined distribution of labware and modules. All labware needs to conform to the Society for Biomolecular Screening (SBS) microplate standard. Consequently, we created a hardware assembly (see **Figure 1A**) that fits on two slots, and which contains three parts: a custom platform in which solid media plates are placed during the entire screening, an image acquisition device and a master computer in charge of connecting the whole set and synchronizing its operation.

**Figure 1.**
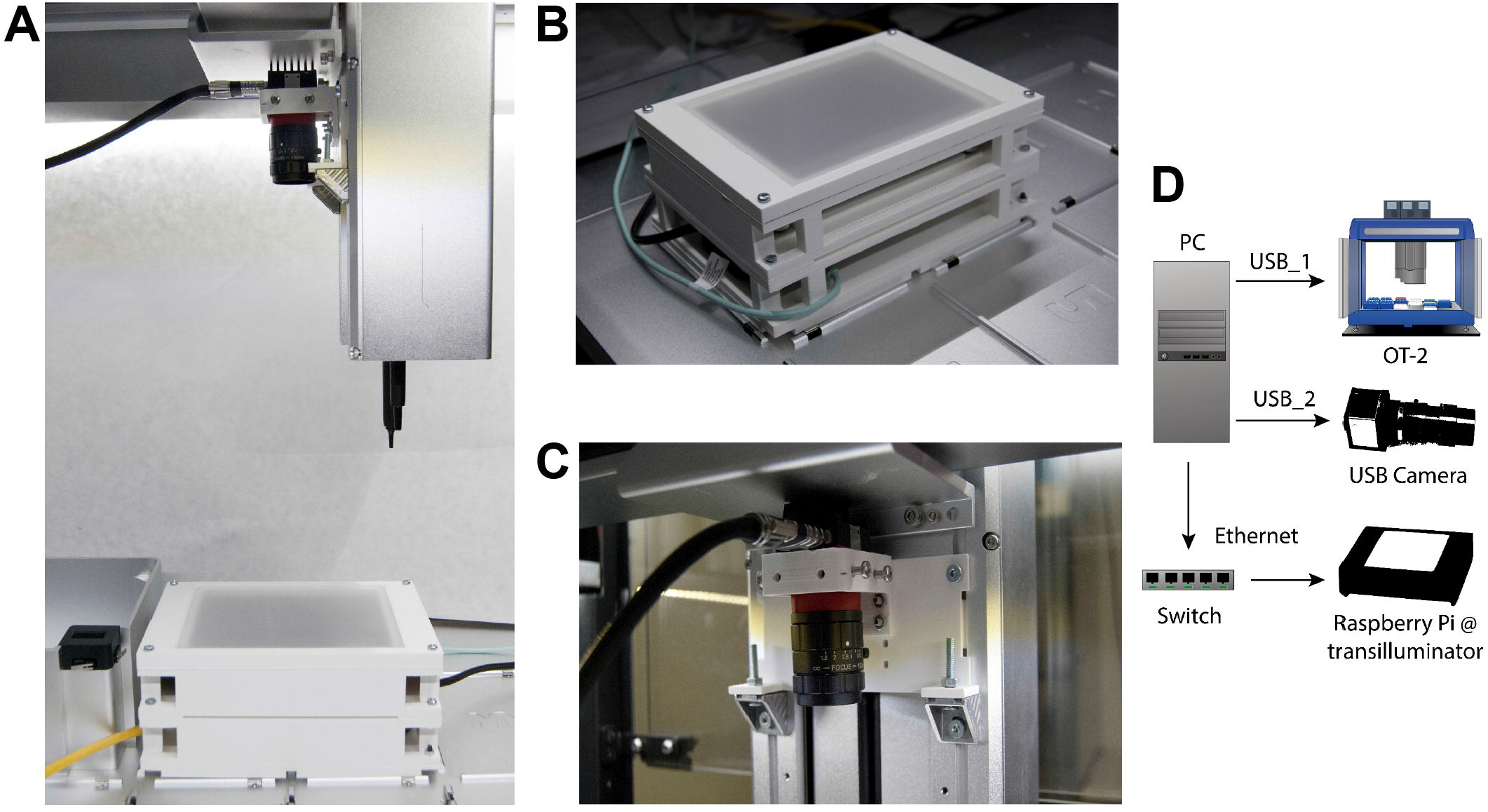
COPICK assembly mounted onto an Opentrons OT-2 liquid handler. (A) The additional hardware consists of (B) a custom-made transilluminator and (C) a USB-camera mounted onto the OT-2 robotic arm (over a P20 pipette) with a 3D-printed scaffold connected to a (D) Master PC acting as a multiport connection Hub.

First, a transilluminator (**Figure 1B**) was assembled using an RGB led panel controlled by a shielded Raspberry Pi 4B. The labware was designed in vertical orientation to allow the use of circular Petri Dishes up to 89 mm. A 3D-printed structure holds the electronic RGB panel and a translucent plastic plate that acts as top cover to support solid media plates and diffuses emitted light during image acquisition. The Raspberry Pi was loaded with Python scripts to switch light remotely using SSH protocol. Image capturing capability was provided by an Alvium 1800 U-500c USB camera with fixed focal distance (Allied Vision) fitted onto the OT-2 pipette arm using a drill-free 3D printed scaffold (**Figure 1C**). The master computer was connected to each part of the assembly to control the execution of the picking protocol: the OT-2 robot and the camera were connected via USB, and the transilluminator was connected via Ethernet using a user-defined local network supported by a network switch (**Figure 1D**).

### 2.2 Inference model description

Inferring the position of colonies in solid plates required the support of an image analysis software to process the pictures gathered by the imaging hardware and identify the pixel regions showing actual real colonies. Although traditional image segmentation algorithms have shown good performance in this context, they are prone to generate detection artifacts under uneven illumination conditions (Huang et al., 2005; Dey, 2019). Their performance is thus limited to a set of particular conditions (i.e. color of the solid media composition, morphology of colonies, color and intensity of light), and any change in those would require a time-consuming calibration. Colony screening should be ideally performed by a more flexible algorithm able to obtain good performance with minimal code modification, which would benefit a broader community of users. In an attempt to overcome such constrain, we chose to integrate a deep-learning algorithm because of their robust prediction capabilities under different image scenarios and its native flexibility to learn arbitrarily complex image patterns (Li et al., 2017; Ha and Jeong, 2021). Concretely, we used Detectron 2, a neural network-based object detection software created by Facebook development team (Wu, 2019). Detectron 2 has a heavily dense layer architecture involving multi-scale feature map extraction, region proposal network and ROI head evaluation that makes it a robust multi-purpose image detection model (Lin et al., 2016; Ren et al., 2017). Among the available set of architectures, this work implemented PanopticFPN, which was specially designed to train panoptic segmentation models (He et al., 2020). Panoptic segmentation is an extended image recognition task, as it simultaneously combines semantic and instance segmentation (Kirillov et al., 2018). Its output is a pixelmap containing not only the class label of each pixel in the target image but also the detected and segmented objects instances. This essentially generates a global and more complete inference about the analyzed image, including both the target object to identify and their related context. This feature, combined with the native robustness of deep neural networks against noise (generated during image acquisition due to experimental fluctuations in light), allows producing better inference results compared with traditional image processing segmentation at the cost of a more complex training dataset. An extensive dataset with annotated images of real Petri Dishes containing microbial colonies was fed to train the Detectron 2 model. Dataset preparation and annotation is described next.

### 2.3 Dataset preparation

Detectron 2 is a machine-learning algorithm that requires a training process using a custom dataset of images containing the ground truth to model, that is, the empirical objects to predict. In the chosen format (panoptic prediction), the dataset must provide segmentation data containing not only objects of interest (colonies), but also about every region present in every image. In this work, we decided to implement a colony prediction model to detect *P. putida* and *E. coli* colonies grown on commercial Petri Dishes of different diameters (outer diameter of 85-89 mm) filled with two different media: LB-agar and M9 salts-Agar (Maniatis et al., 1982). To create the dataset (see **Figure 2**), we first obtained a high-quality set of images. A sample of 200 images of real agar plates containing a total number of 18660 colonies was gathered using a reflex camera with a macro objective (Nikon D60 with AF-S Micro NIKKOR 60 mm f/2.8G ED lens) assembled into a copy stand (Gelprinter). Illumination of images was optimized by using a commercial circular white LED lamp (inspire Manoa) coupled with an in-house diffusor filter composed by translucent layers of Polyethylene terephthalate (PETG). Each image was annotated into an info database in *.csv format containing data about plate preparation (i.e. number of colonies, type of medium, etc.) that was used during image segmentation step. Next, we used augmentation techniques to multiply the training data available to feed the model (Shorten and Khoshgoftaar, 2019).

**Figure 2.**
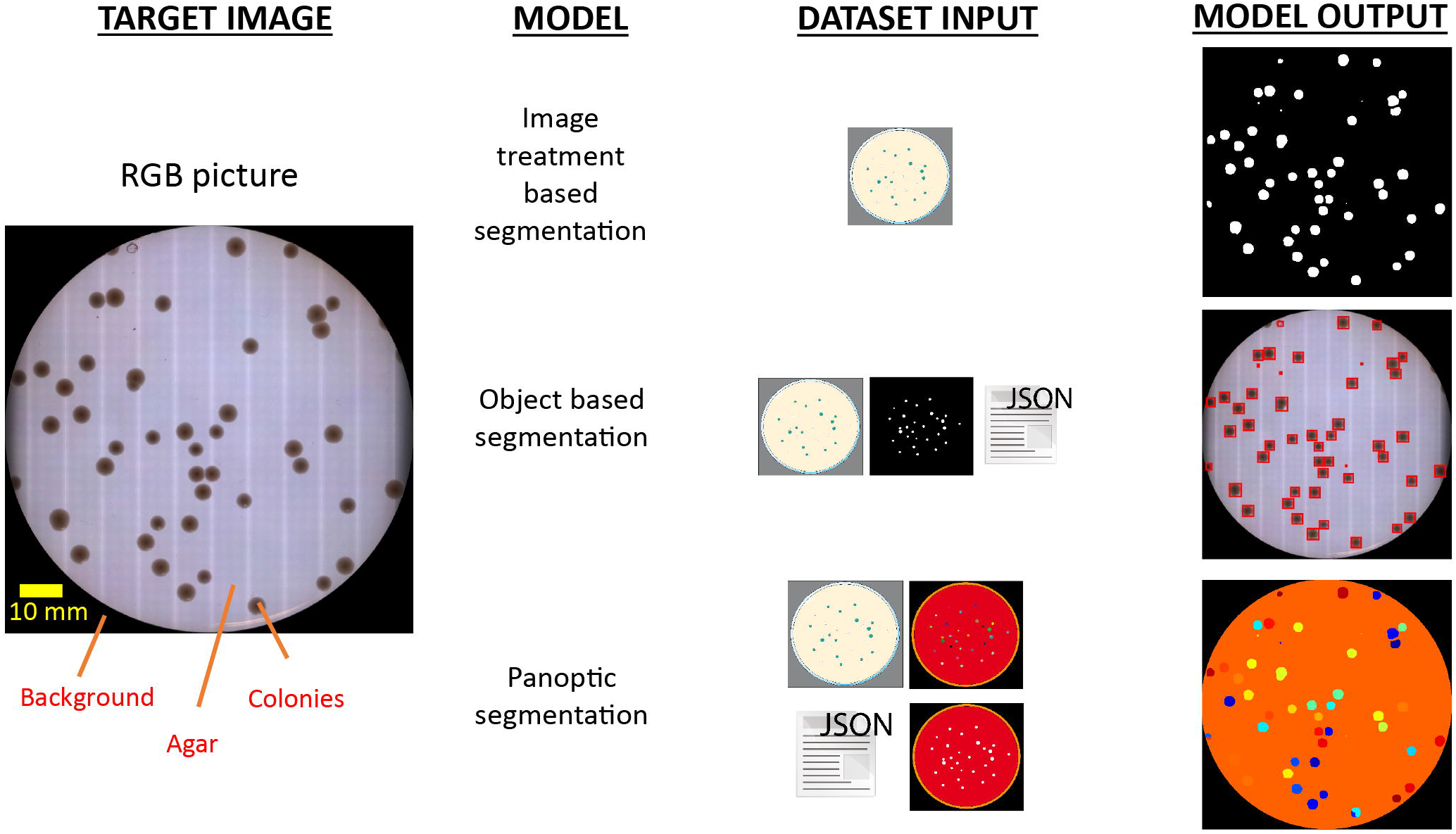
Informative tradeoff between the required input data and delivered output for different segmentation techniques to discriminate colony presence on agar plates. Identification of colonies has been historically addressed by image treatment segmentation techniques based on detecting local changes of pixel intensity values in the image (up). As a consequence, this approach demands tightly controlled imaging acquisition conditions to ensure robust results. The use of object segmentation models (middle) allows to relax this strict constrain by applying a learning stage in which a dedicated dataset with object-level annotations is required. These models perform object discrimination by detecting specific geometric and pixel intensity patterns of target bodies, and typically output the regions of interest (ROI) enclosing identified objects. However, COPICK makes use of a panoptic segmentation model (bottom), which performs a class-based inference at pixel- and segment-level on input images. This is achieved by feeding the algorithm with raw images, panoptic and semantic masks and detailed annotations about the different segments composing each image.

From each image, we obtained 15 derivatives by applying operations of rotation and mirroring. The final dataset totals 3000 images and 279900 colonies. Once the expanded image dataset was generated, we applied a customized segmentation algorithm to generate the pixel mapping of each image, which was required for training panoptic inference. **Figure 3** shows a general scheme illustrating the image processing applied to each image of the augmented dataset.

**Figure 3.**
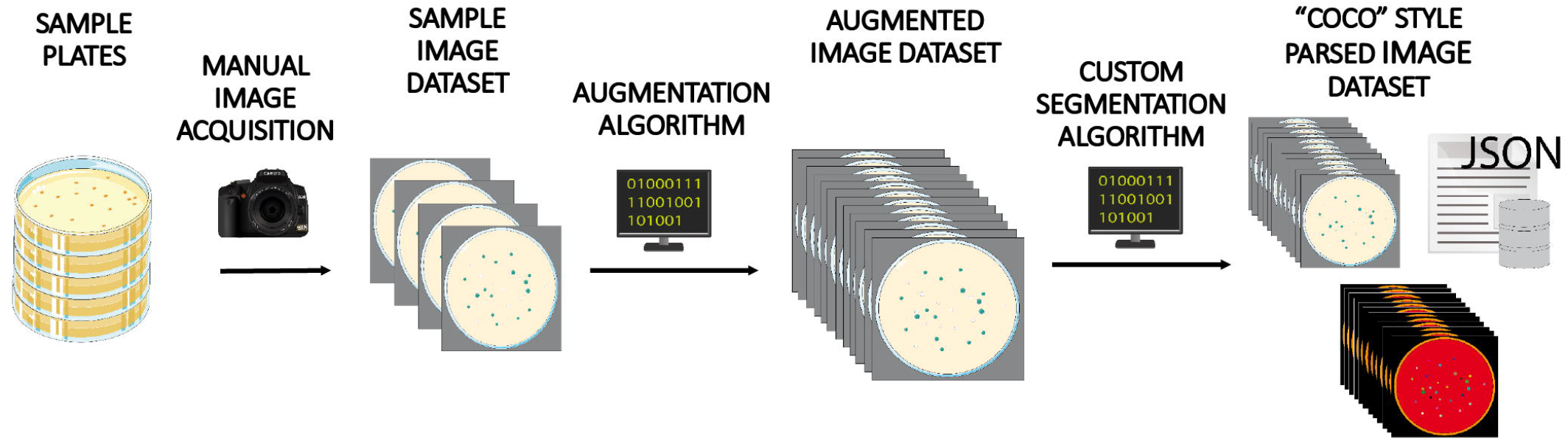
Dataset creation workflow. A batch of 200 Petri dishes containing *E. coli* and *P. putida* colonies grown on different media were manually photographed and annotated. This image set was next augmented 15-fold by applying different image transformations. Finally, each image was processed using a custom-made image treatment algorithm to infer panoptic segmentation and create and parse associated annotations and masks in COCO style format, one of the file structures accepted as valid input by Detectron 2 model.

Briefly, raw input images were grayscaled and reduced in size by cropping the Petri dish bounding box. A masking process divided the plate in two parts that were treated separately: the inner part was binarized using a combination of Gaussian blurring and adaptive Gaussian thresholding method (Shapiro and Stockman, 2001; Roy et al., 2014), and further treated with a 2D Sklansky algorithm (Sklansky, 1982) to obtain the convex hull of binarized colonies. Petri dish border image was treated by applying a speckle noise addition step to ease the removal of plastic borders using a canny edge detection operator (Canny, 1986), and a final sequential filtering stage using erosion, hole filling and top hat operations (Shapiro and Stockman, 2001). Parameters of algorithms were manually fitted to obtain estimations in colony counting with errors below 5% of annotated values in the info database. Panoptic segmentation data was created using discrete categories labeled as “colonies” and “background”. Additionally, “colonies” objects were sublabeled according to the number of actual colonies: “1 colony”, “2 colonies”, etc. Regions categorized as “background” were sublabeled into “out of plate” (raw background) and “0 colonies” (regions within the plate with no colonies: agar) categories. Finally the augmented image repository and its complementary panoptic segmentation info were parsed and stored into a set of annotated *.json and *.png files following the data structure given by COCO dataset style for PanopticFPN models (Lin et al., 2014).

### 2.4 Training, Evaluation and optimization of inference model

Figure 4. shows a description of the workflow followed to train and optimize the inference model for colony detection. Input dataset was split into a training and validation subsets containing 75% and 25% of images respectively. Prior to training, it was necessary to set up the hyperparameters that control the behavior of the model related to neural network performance (i.e., the architecture of data processing layers, the duration of the training process or the learning rate, etc.). Once chosen, the training algorithm ran for a variable duration between 1 and 24 hours. At the end of the run, the resulting model was stored. Panoptic segmentation inference for any input image takes just a few seconds. The predicting power of the trained model was monitored after each training run. During this stage, a set of metric scores were computed based on the inference performed on the validation subset. In this work we selected two proxies assessing panoptic prediction performance: Panoptic Quality (PQ) and Segmentation Quality (SQ) (Kirillov et al., 2018). Where PQ informs about the estimated performance of the model to properly segment and classify all the present category types and objects in the images, SQ focuses on evaluating the capacity of the model to properly segment the morphology of detected objects within the images. The value of these scores was dependent directly on the hyperparameter choice, thus it was necessary to select them properly.

In order to choose good values, we used the Optuna optimization package to implement the execution of a Tree-structured Parzen Estimator algorithm combined with a trial pruner using a median stopping rule (Bergstra et al., 2011; Akiba et al., 2019). The optimizer executed recurrent runs of the object function (a complete Detectron 2 training-evaluation session) making tentative guesses to the hyperparameter set given as input, trying to maximize the output performance score. We performed two rounds of optimization with a sample size of at least 50 trials using PQ and SQ separately, storing the output set of weights generated at each trial. Next, we evaluated the inference capability of the whole set of weights by benchmarking their precision on detection (number of detected colonies and size deviation) for a random selection of images in the dataset (see Discussion section for more details). Finally, we analyzed the predictive nature (conservative vs greedy) of the preselected set of weights and chose a balanced triad for operational purposes in real colony detection tasks. The final output of the inference stage was estimated as the consensus at pixel level between the three predictions performed using this set of weights.

**Figure 4.**
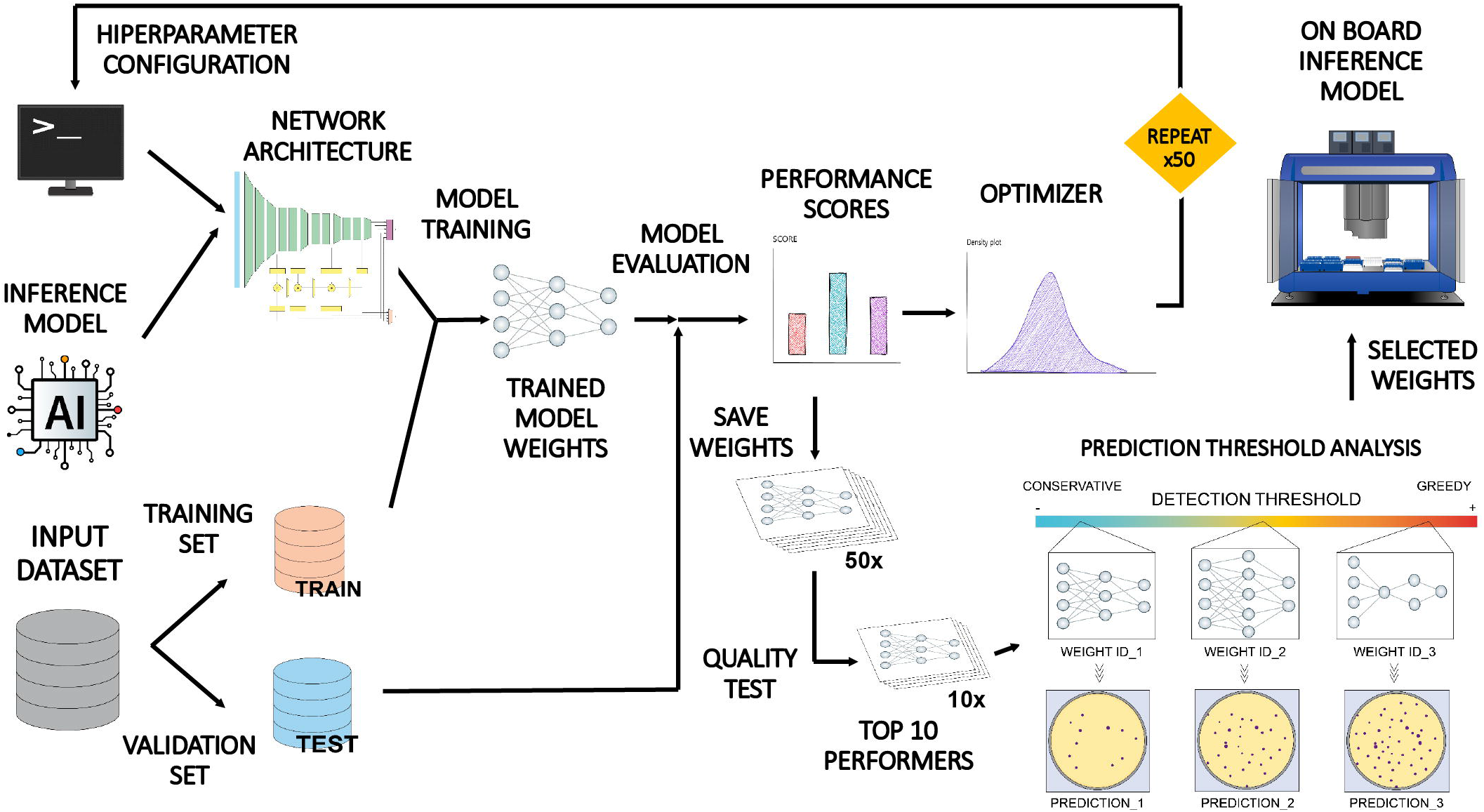
Schematic illustration of the training, validation and optimization workflow applied to COPICK inference model. The Detectron 2 loaded with the PanopticFPN model was configured by assigning tentative values to a set of hyperparameters setting up its architecture. The model was trained and evaluated using two subdivisions of the input dataset. Obtained performance scores (Panoptic Quality PQ and Segmentation Quality SQ) were separately used as proxies to guide a fine-tuning of hyperparameter configuration by using an optimizer algorithm in two independent rounds of > 50 iterations. All resulting model weights were benchmarked using a colony counting and geometry matching test to select top 10 performers. Finally, a prediction threshold analysis was applied to select two final sets of weights yielding a conservative, moderate and greedy predictions that were used to generate the final inference in the robot by consensus.

### 2.5 On-board colony screening methods

Panoptic segmentation inference delivered the outputs of the footprint of detected colonies. This footprint was used to gather visually related properties. In this implementation we analyze three features typically checked in colony screening protocols: colony size, colony color and colony brightness.

Colony size was directly obtained from panoptic prediction as the raw pixel area occupied by every segment contained in colony footprint. Size discrimination in real mm was derived by estimating the pixel to mm scaling factor associated with image acquisition camera. Colony color was retrieved by computing the pixel-average HSV color within the colony footprint. With this information, a color filter was programmed to select or exclude candidate colonies based on a manually selected color range. Additional screening capabilities based on fluorescence emission were achieved by mounting a low-pass filter at desired wavelength and using the light selecting capabilities of the RGB panel. In this case, we mount a GFP filter (500 nm filter, Edmund Optics Hoya Y50) and retrieve two images: a regular one using white light and an additional image switching only blue LEDs of light panel which excite the GFP (maximum intensity around 460 nm). The former was used for colony footprint prediction, where the latter gathered the specific fluorescent emission of colonies.

The screening system was programmed to generate a normalized unitary score of every detected colony based on a user-configurable function. In this work we selected a weighed sum of individual marks attending to the three above mentioned features (size, color, fluorescence) to make available the possibility of using a multicriteria screening capability.

### 2.6 Robotic picking of colonies

To physically perform colony picking, OT-2 must know the exact physical coordinates of each colony to pick in robot coordinate reference system. However, colony coordinates are natively obtained in local pixel reference system of the image captured by the camera. Coordinate mapping between local pixel-based reference to robot-based reference (in mm) involved a set of transformations described below and summarized in **Figure 5**:

**Figure 5.**
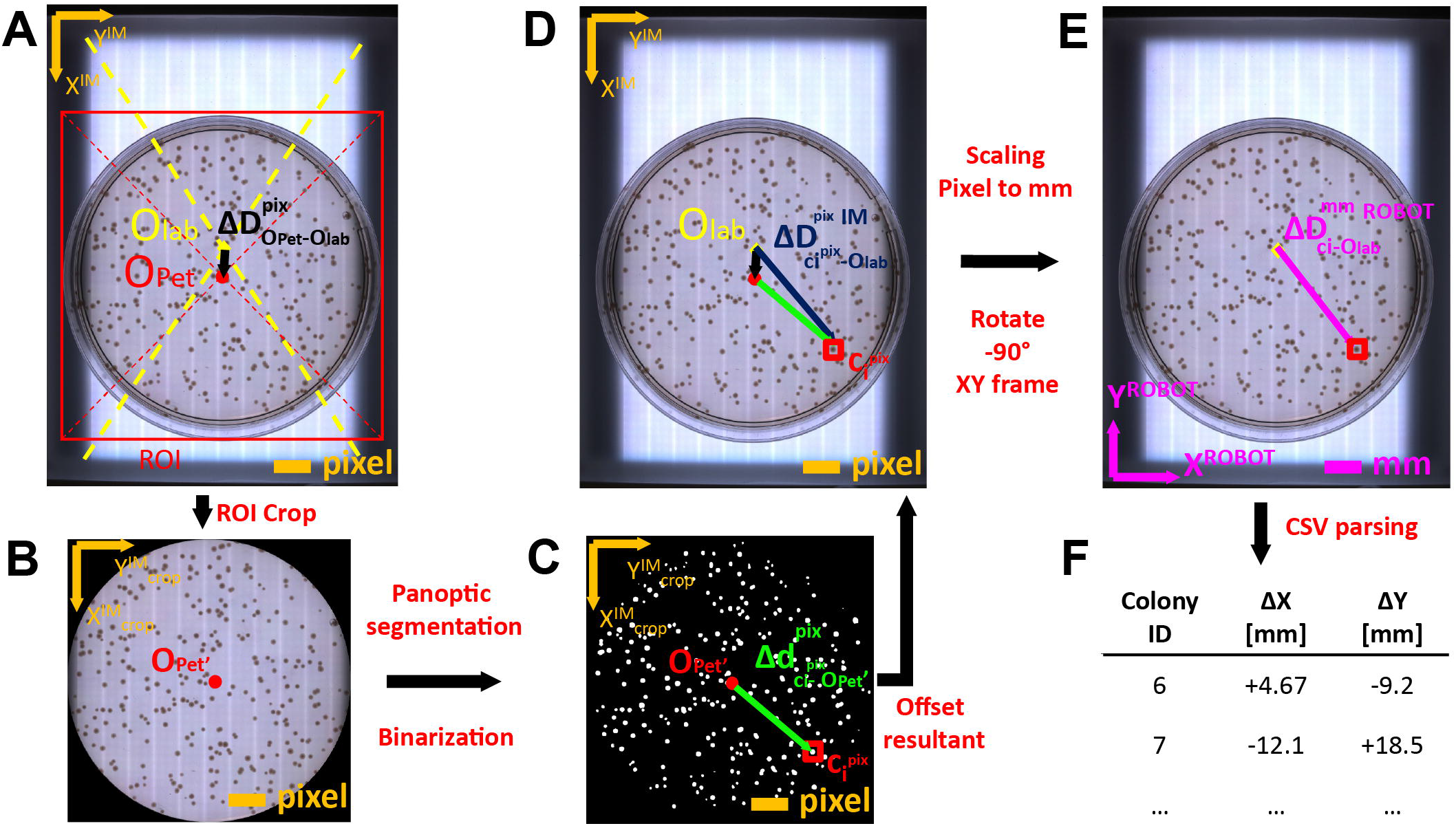
Coordinate mapping of detected colonies from image coordinate frame to Opentrons 2 OT-2 robot frame. (A) Images acquired by USB camera on transilluminator are analyzed to infer the pixel offset (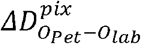, in black) between the center of the labware (O_lab_) and the geometrical center of Petri dish (O_Pet_). Cropping the agar plate region ROI encased by its bounding box and discarding plastic borders allows to obtain (B) a clean image to feed into the inference model. (C) Segmentation and object binarization of detected colonies allows calculating the pixel offset (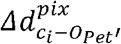, in green) between the colony centroids (c_1_) and the local center of the Petri dish (O_Pet_’) in the crop image. (D) The pixel offset between the colony center and the center of the labware is the vectorial sum of both offsets (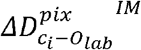, in blue). (E) Pixel offset vector is scaled to mm using the camera pixel-mm conversion factor and the coordinates are rotated 90° anti-clockwise to match robot reference coordinate system (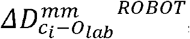, in pink). (F) Finally, the process is repeated for each detected colony and the computed offsets are annotated into a *.csv file that will be uploaded to Opentrons to perform physical picking.

1. Gather an overview image of the transilluminator area with a Petri Dish on it. Find the center of the labware (Olab must be previously calibrated, see Supplementary Material).
2. Find the bounding box containing the Petri Dish and retrieve its center OPet in overview photo pixel coordinates.
3. Compute the offset between center of transilluminator labware (previously calibrated, O_lab_) and Petri Dish (O_Pet_) center in pixel coordinates 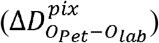.

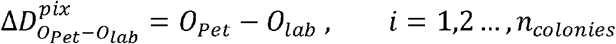
4. Infer panoptic segmentation by feeding a cropped image containing the isolated bounding box to the trained model.
5. Binarize panoptic segmentation (zeroing agar and background coordinates) to detect object centroid positions (colony coordinates 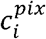) and store Petri-Dish center (OPet*’*) in local crop pixel coordinates.
6. Compute colony position offset between colony centroids and plate center in pixel coordinates.

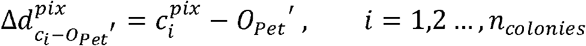
7. Rescale offsets in row (x) and columns (y) from pixel coordinates to mm using scaling function f = [*f*_*x*_,*f*_*x*_]. Note that each axis has its own transformation (must be calibrated first, see Supplementary Material).

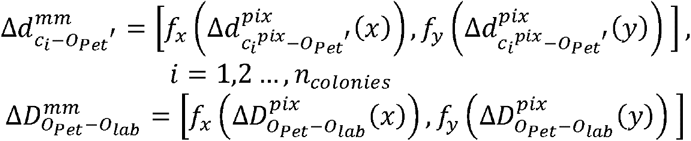
8. Compute offsets (scaled in pixels) from colony positions (ci) to center of labware (Olab) by vector addition in image coordinate system.

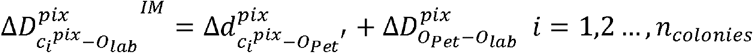
9. Compute offsets (scaled in mm) from colony positions (ci) to center of labware (OPet) by vector addition in image coordinate system.

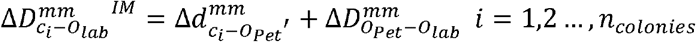
10. Apply a 90° anticlockwise rotation to shift coordinates gathered image coordinate system to robot coordinate system.

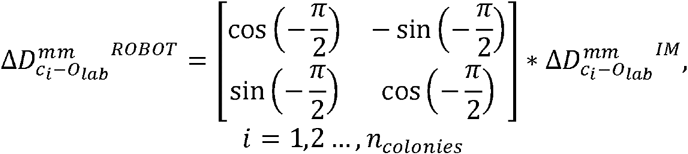

Colony coordinates were finally annotated into a *.csv file (after applying selection filter) and uploaded to OT-2 robot for colony picking execution.

### 2.7 Automated workflow and Orchestrator software

After assembling the hardware parts and training the inference model, it was necessary to define a workflow addressing the different actions to execute by the hardware (OT-2 robot, transilluminator, camera) and software (inference model) during colony screening and picking. This set of instructions, illustrated in **Figure 6**, can be summarized as follows:

**Figure 6.**
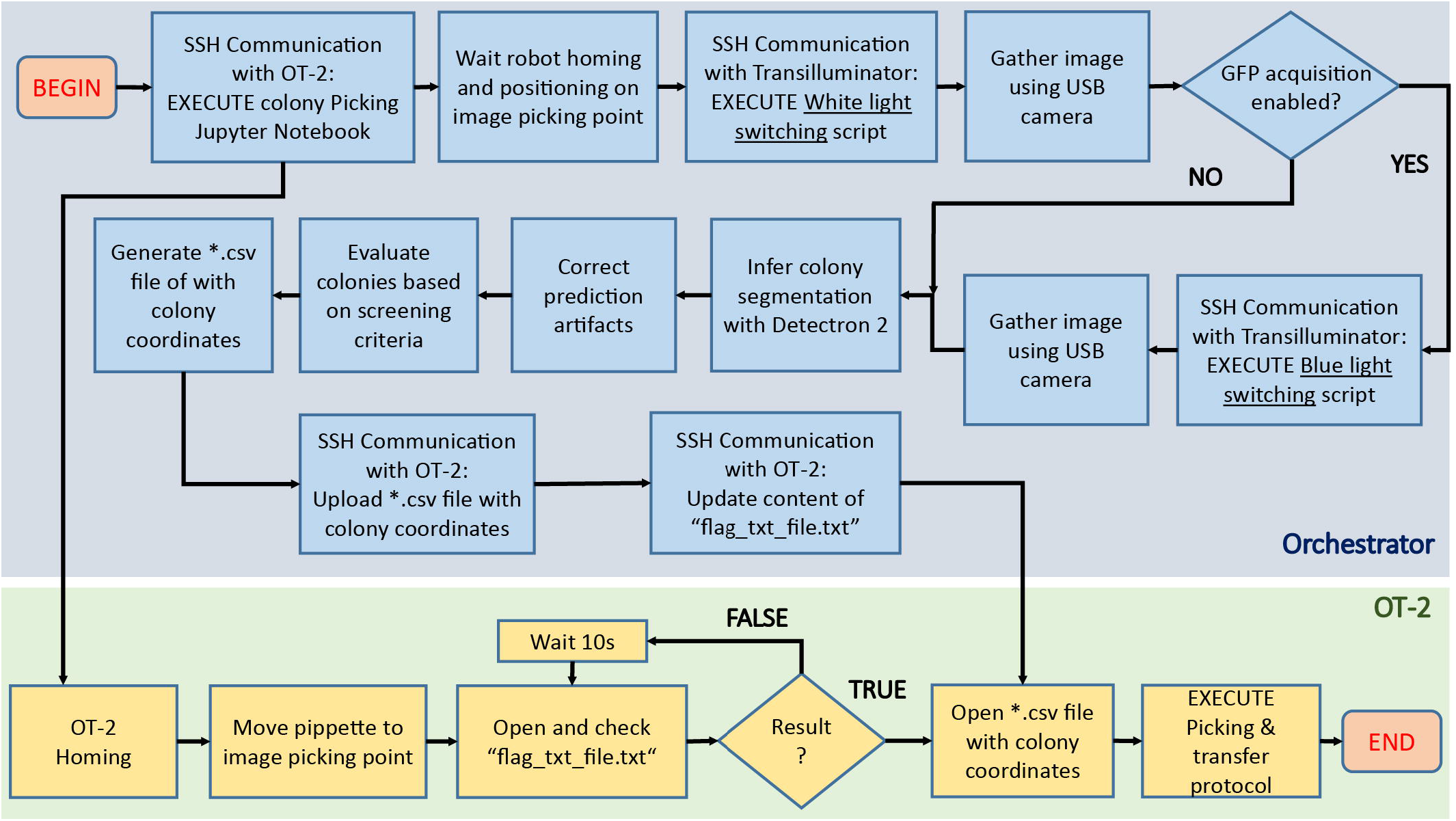
Algorithmic pipeline executed by COPICK package to execute a colony screening and picking protocol. After placing the agar plate onto the transilluminator, the orchestrator software (blue area, up) sends an SSH command to the OT-2 server to execute a custom Jupyter notebook containing the colony picking instructions. The robot (green area, bottom) initializes the protocol by first homing and next moving the pipette to a preset image acquisition position. A flag-dependent loop halts further execution of picking protocol in OT-2, allowing the orchestrator to start the image acquisition process: an SSH command is sent to the Raspberry PI hosted in the transilluminator to switch the activation of white light, and then gather an image using the USB camera. If a GFP based screening is required, the process is repeated with a blue light instead. After images are gathered, they are processed and fed to Detectron 2 model to obtain a preliminary colony segmentation, which is further refined using image treatment techniques. Screening of detected colonies is implemented next by performing a score-based selection using the user-defined criteria (size, color, fluorescence). Coordinate mapping is computed to obtain the positions of colonies in robot coordinates, and its result parsed into a *.csv file uploaded to OT-2 via SSH scp protocol. Finally, the content of the auxiliary file (“flag_txt_file.txt”) is updated to raise the loop flag and unlock OT-2 workflow: the robot accesses the file with the colony coordinate list and start the physical picking process.

1. Place camera in acquisition point
2. Activate transilluminator
3. Image acquisition
4. Generate prediction with Detectron 2
5. Correct prediction artifacts
6. Filter colonies based on user-defined criteria
7. Create *.csv file with physical coordinates of target colonies
8. Perform physical picking

To synchronize the execution of this workflow, a custom-made orchestrator software was programmed to interface operations performed in peripherals with the main workflow executed by the PC. Communication between PC, transilluminator and OT-2 was generated via SSH.

Following the mentioned scheme, the robot must first place the camera in position and wait for *.csv file containing positions of colonies to proceed with picking. As OT-2 does not provide possibilities to remotely control the robotic arm using unitary instruction commands, we implemented a naïve solution based on uploading an OT-2 protocol containing a file-content based flagging condition. This script is executed at the beginning of any colony picking protocol via SSH by the orchestrator. The script sends instructions to OT-2 to allocate the camera in position, and then execute a while loop in which the content of an auxiliary .*txt file is recurrently inspected. The picking protocol cannot continue unless such content is updated with the proper keyword. While OT-2 is looping, the orchestrator software continues with the execution of the rest of the workflow until coordinate file is generated and uploaded to OT-2. The content of the auxiliary file is then updated, allowing the OT-2 script protocol to reach the loop exit condition and order the robot to first retrieve the uploaded coordinates from its own memory and finally proceed with the picking process. Details about each step of this workflow can be found in Supplementary Material, code repository readmes and code commentaries.

### 2.8 Benchmark tests

Three different tests were performed to evaluate the reliability of COPICK as a colony picker. First, we measured the accuracy and precision of colony detection and picking without any selection criteria. A total number of 7446 colonies distributed in 44 Petri Dish plates containing *E. coli* and *P. putida* colonies within the range 12-453 were gathered and screened. For each plate, we annotated counts of detected and picked colonies, detected and picked artifacts (noise), not pickable colonies (colonies next to the plastic border), not detected colonies, single colonies picked multiple times and colony groups detected as single colonies. Evaluation metrics (raw performance, accuracy, misclassification rate, sensitivity and precision) were computed after deriving aggregated data of true positive (TP), False positive (FP) and false negative (FN). Details about metrics and data processing are explained in Supplementary material and accessible in the Supplementary Table. Also, the benchmark image set containing gathered pictures of the plates and the inferred panoptic predictions are available in repository folders.

Next, we tested the screening capacity of COPICK based on color measurement. To do that, we prepared LB-agar plates supplemented with X-gal at 50 µg/ml, and inoculated them at proper dilution factor with a 50-50% mixture of *E. coli* DH5α carrying two pSEVA plasmids: pSEVA637 (colonies exhibit a white color) and pSEVA 6819[gD] (colonies exhibit a blue color in presence of X-gal) (Martínez-García et al., 2023). Based on a predefined target RGB color (corresponding to an arbitrary selection of blueish color displayed by pSEVA 6819[gD] strain), we programmed the selection of 64 clones exhibiting the most dissimilar color respect to the given reference in descending order (column-wise order), transferring them into a single-well plate (Nunc Omnitray, ThermoFisher Scientific).

To further check COPICK selection capabilities, an additional validation test was prepared to benchmark its proficiency during a fluorescence based screening. *P. putida* TA245 (Aparicio et al., 2020) and *P. putida* KT-GFP (Espeso et al., 2016) strains were plated together in order to check whether the implementation could discriminate both genotypes based on the different levels of GFP-based fluorescence that their colonies display (the former was as negative control of GFP signal, compared with the later showing a strong fluorescent signal). Respective overnight cultures of both strains grown in LB-Lennox broth were mixed using a 50:50 ratio, and plated on a LB-agar plate at required dilution to incubate overnight at 30°C. Colony screening consisted on selecting the 30 most fluorescent clones in descending order (column-wise order) of fluorescent brightness, transferring them into a single-well plate.

## 3 Results

The performance of the proposed technical adaptation was benchmarked by executing colony picking protocols under three different screening scenarios: raw colony picking (no screening) and phenotype cherry-picking based on visible color and fluorescent reporter.

The first test was performed to measure the degree of precision delivered by COPICK adaptation when picking colonies. **Figure 7A** shows a summary of the obtained results. Generally, all metrics are satisfactory, with all values above 70%. Obtained raw performance over total screened colonies scores 73%, a number that raises up to 82% if only pickable colonies are considered. By “pickable colonies” we mean those colonies which are not in close contact with the border of the Petri Dish, in such a way that an attempt of the robot to access the colony would have a large risk of hitting the Petri plate. This security margin was applied to avoid causing mechanical ruptures of the plastic dish or bending the tip. The sensitivity level of 92% indicates low levels of noise-related issues, whereas a 73% of accuracy in combination with a precision of 78% points towards the existence of issues counting colonies inside groups that diminish the overall effectiveness.

**Figure 7.**
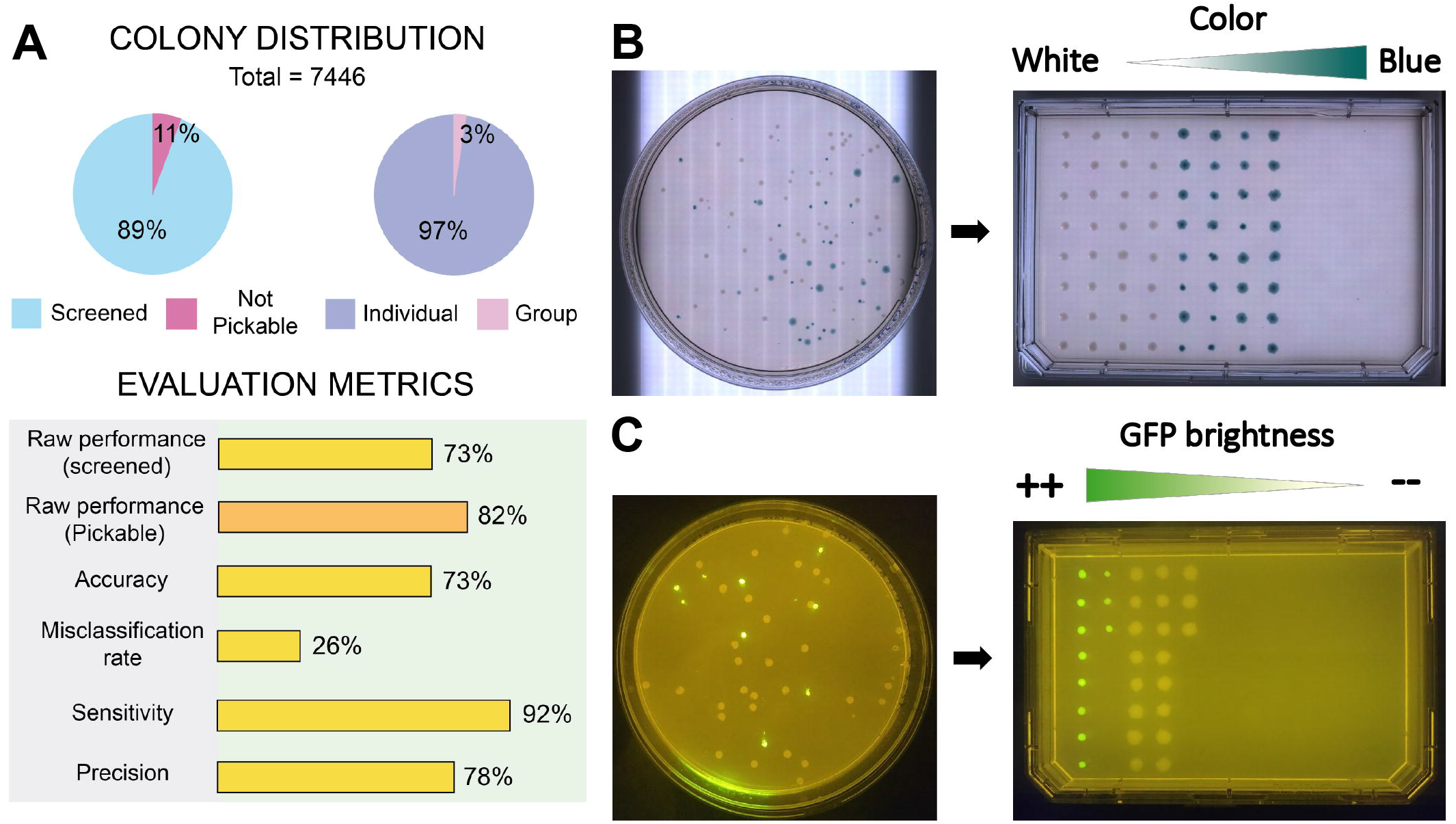
Results of benchmark tests performed to COPICK implementation. (A) Sample distribution of screened colonies and quality metrics obtained during raw picking test. (B) Naked eye and (C) fluorescence images showing the output obtained during validation assays of phenotype screening based on visible color and GFP reporter.

Positive results obtained in colony filtering tests based on color and fluorescence indicate that COPICK can process images to successfully screen colonies based on these phenotypic traits. **Figure 7B** shows a selection of colonies showing a mixture of white and blue phenotypes based on a dissimilarity criteria respect to a reference color (blue). COPICK executes a picking protocol where those colonies with best matching color (blue) are picked last (picking order is set by column), whereas the most different (white) are picked first. The output of the fluorescence-based screening under a blue light transilluminator (**Figure 7C)** exhibits similar results: colonies are picked following a descending order in average intensity. Both protocols can easily be adapted to select and pick the colonies as required.

## 4 Discussion

Despite its interesting performance, COPICK has some limitations found at hardware and software level that are worth mentioning. Regarding the software, results delivered by inference model suggest a technical issue during the training stage. By visualizing the classification results delivered by Detectron 2, it can be observed a very poorly object classification performance: the number of colonies detected as “objects” roughly exceed the 50%. We hypothesize that the poor yield exhibited by the model identifying colony objects might be caused by a combination of two issues. First, an error in the dataset input format could be interpreted by Detectron 2 as an unrecognized data, and thus ignored. This is somehow in agreement with the results obtained during hyperparameter optimization process: we realized that, on average, larger values of PQ and SQ delivered better inference results. However, we oppositely found that those sets of weights obtaining the largest scores were not the best predictors. Next, more images could be required to improve colony prediction, as the overall pixel-area occupied by colonies in the current dataset might not be enough to produce a robust panoptic segmentation (microbial colonies are relatively sparse tiny regions inside images). This would be in agreement with the fact that the model performs an excellent segmentation of agar footprint, and with the result of delivered output: where the model cannot identify a colony and the pixel is not agar, the segment is assigned with a zero-value label. Interestingly this result delivers a *de facto* accurate prediction: the inference of microbial colonies is indirectly obtained by combining the segments positively predicted as colonies with the subset of non-recognized pixel regions.

A second point to consider is the cost of using a panoptic segmentation model. In this work we had to develop a custom-made script based on image treatment algorithms to obtain the required segmentation and annotations of dataset images, at a cost of investing time and resources. At present time (early 2023), there is still a lack of open-source tools on the web to label and create datasets of images using a panoptic segmentation format in a straightforward fashion. Any user interested on using COPICK to identify other type of colonies will need to create its own segmentation script or modify the one included in this package to successfully create a good dataset for training. As a remark, using a panoptic dataset allowed us to obtain a remarkable picking score and to generate a robust portable model to be used into OT-2 transilluminator device (despite the fact optic and illumination settings were worse than those use for database creation).

Another limitation observed when analyzing the results of the presented inference model was the correct detection and interpretation of colony groups. With the presented dataset, COPICK inference model was unable to learn how to split properly individual colonies forming part of colony groups. The reasons of such limitation could be the relative low number of colony groups (respect to individual colonies) presented to the inference model during the training stage. To fix this issue, we implemented a postreatment stage during orchestrator workflow execution in which grayscale regions matching inferenced objects were further analyzed using a local maximum search algorithm to find multicolony patterns. This approach succeeds on dividing most of segments composed by more than one colony at a cost of introducing spurious divisions of single colony footprints colonies at low event frequency. Although we attempted to remove this source of error using different approaches (such as object filtering criteria based on aspect ratio or circular pattern search based on Hough transform), we did not no succeed on it. In fact, a general solution to this problem could be hard to achieve for those cases in which colonies exhibit different growth rates or a non-circular growth pattern. As a suggestion, a trivial solution to this issue would be the development of a graphic interface to allow the user discard or include elements of the generated inference.

When talking about the hardware, we found that OT-2 stepper motors in charge of positioning the robotic arm in place introduce an intrinsic error when picking colonies. We speculate that OT-2 NEMA motors were not assembled to work with such degree of spatial accuracy, but to target positions in (open) well plates. As a consequence, picking small colonies is a task biased not only by the positional error introduced by the inference model, but also enhanced by the addition of this error source. In fact, we observe substantial picking error when trying to transfer colonies with diameters below 1 mm. This issue, however, can be easily overcome by screening plates with larger colony sizes. Additionally, we detected that OT-2 was causing inconsistent picking patterns due to the generation of vibrations in the labware deck, associated to default settings of robotic pipette movements (displacement and tip picking). To solve this issue, we designed an adaptor to tightly hold Petri Dishes of different diameters on place (can be found in package repository), although this solution implied a less comfortable manipulation.

Finally, we realized that performing a good selection of colonies based on fluorescence and color was dependent of achieving minimal quality background levels of brightness and contrast. Concretely we identified issues related to this observation close to the right and left borders of the RGB panel. Although COPICK inference model was able to properly discriminate colony footprints on these problematic areas, screening filters sporadically failed on evaluating their correct color/brightness values. A larger RGB panel should fix the problem, but occupying more slots of the OT-2 deck.

A similar situation was observed when fluorescent brightness based discrimination was evaluated: succeeding was only possible when using strains exhibiting a medium-high light intensity signal. The long-pass filter assembled during the fluorescence based cherry-picking validation test was not able to provide enough dynamic range to discriminate those colonies exhibiting a poor GFP signal. Furthermore, other fluorophores excitated at similar wavelengths and broadly used to discriminate strains or genetic constructs (i.e. mCherry) were not discriminated either. Acquiring a more specific and sensitive filter should fix this issue.

## Supporting information

Picking test data table

## 5 Conclusions

Benchmark tests performed under three different screening scenarios (raw colony picking, color based and fluorescence-based cherry-picking) validates the utility of COPICK as a reliable technical solution to provide a colony picking solution into an OT-2. The setup can be easily deployed by connecting an OT-2 robot to an open-source hardware assembly composed of a custom 3D-printed labware, a USB camera and a computer able to run the trained inference model. Its minimal implementation cost (in terms of time, expertise and resources), compared to commercial solutions for picking operations, converts this solution in an attractive alternative to introduce automated colony screening workflows in microbiology laboratories.

## 7 Permission to reuse and Copyright

Provided scripts in <u>COPICK package</u> are under **MIT license**, and thus freely available for distribution, modification and use. <u>COPICK dataset</u> is under **CC BY 4.0** license, and thus freely available for distribution, modification and use respecting attribution right.

## 8 Conflict of Interest

The authors declare that the research was conducted in the absence of any commercial or financial relationships that could be construed as a potential conflict of interest.

## 9 Author Contributions

DR and JN contributed to conception and design of the study. DR and IO programmed the Python scripts for database creation and parsing. IO created and structured the image database, performed the training, validation and optimization of the inference model. DR, IO and PY designed and tested CAD models of image acquisition system. PY programmed the acquisition scripts for USB camera. DR designed and programmed the transilluminator labware. DR programmed orchestrator software and OT-2 picking scripts. IO and AGL performed the hardware benchmark tests. DR wrote the first draft of the manuscript. All authors contributed to manuscript revision, read, and approved the submitted version.

## 10 Funding

This work was supported by the European Union’s Horizon 2020 Research and Innovation Programme under Grant agreements no. 814650 (SynBio4Flav), 870294 (MixUp), and 101060625 (NYMPHE). Funding was likewise provided by the Spanish Ministry of Science and Innovation under “Severo Ochoa” Programme for Centers of Excellence in R&D, grant SEV-2017-0712 (AEI/10.13039/501100011033) and the RobExplode project: PID2019-108458RB□I00 (AEI /10.13039/501100011033). JN and AG acknowledge the financial support of CSIC’s Interdisciplinary Platform for Sustainable Plastics towards a Circular Economy+(PTI-SusPlast+). IO and AGL were funded by GARANTIA JUVENIL CAM 2020 program of the Comunidad de Madrid – European Structural and Investment Funds –(FSE, FECER) through grants PEJ-2020-AI/BMD-18724 and PEJ-2020-AI/BIO-18028, respectively.

## 11 Acknowledgments

We thank Ana Anhel, Blas Blázquez and Juan Poyatos for helpful discussion.

## 12 Supplementary Material

The provided supplementary material package includes a supporting information document and a supplementary excel file summarizing the performance statistics. Image collection used for validation can be found in Github repository indicated below.

## 13 Data Availability Statement

COPICK package containing python scripts, CAD designs and user guides can be found in the following GitHub repository https://github.com/sysbio-cnb/COPICK. Dataset containing images of Petri Dishes gathered to train and validate Detectron 2 based inference model can be found in the following URL: https://sysbiol.cnb.csic.es/EMBBT/COPICK_DATASET.zip.

## Notes

https://github.com/sysbio-cnb/COPICK

https://sysbiol.cnb.csic.es/EMBBT/COPICK_DATASET.zip

### Competing Interest Statement

The authors have declared no competing interest.

