## Supplementary information for "Technical upgrade of an open-source liquid handler to support bacterial colony screening"

Supplementary Material

### 1 Orchestrator software description

The orchestrator software is a python written script that regulates and synchronizes the execution of the different hardware and software parts of COPICK technical solution. It contains a set of variables and functions that can be modified by the user according to their needs. Details are given following the structure of the script. These steps match with the algorithm description depicted in Figure 6 of the main manuscript:

a) Flag_txt_file initiation and upload. Flag file in charge of halting picking of OT-2 is created and set to value “NO”. The file is uploaded to OT-2 via SSH.
b) Execution of picking colony protocol loaded in OT-2 via SSH command.
c) Wait for robot homing and positioning of camera in place. This process lasts around 60 seconds.
d) Execute image gathering script. The script opens an SSH connection with the Raspberry Pi and executes the white light switching script loaded in it. We provide different versions of the switching script with different time durations. By default, we set this duration to 3 seconds. Then, an image is taken using the camera settings defined at the beginning of the script. Note that camera settings (light exposition) and hole aperture must be calibrated to obtain an image with enough contrast. The image is first rotated 90º to obtain a front view of the plate, then named according to the day and time acquisition and saved. If gfp is activated, then repeat the same process but using the blue light switching program. Note that gfp images must be grabbed using the blue light filter and in a dark room.
e) Execute colony detection algorithm. White light image is first cropped to reduce the number of pixels. To do that, image is first grayscaled, then the edges of the image are computed using a canny algorithm to further perform a circle detection based on Hough transform and peak local maxima (using 5 possible peaks) algorithms (Shapiro and Stockman, 2001) with a radius chosen for the size of the used Petri dishes. Note that these may vary depending of the commercial brand used, so it is required to check this parameter to obtain a good circle detection. The center of the Petri dish will be the average of the provided center coordinates. With the center and the radius, image is cropped using a squared bounding box with length equals to the diameter of the circle, and the pixels out of the plate are dropped to zero by using a circular mask with the radius of the detected circle. The coordinate of the circle center is returned for further use, and the cropped image is stored in a new file.

Next, the cropped photo is passed to feed the inference model for colony detection. Inference will be applied three times using three different sets of weights with different prediction behaviors. These adjectives refer to the tendency of each model to generate predicted segments whose pixel-based areas are reduced (conservative), average (moderate) or large (greedy) in value compared to real colony footprint over which inference is being performed For each detection, Detectron is loaded with the respective weight file and the network evaluated. Output panoptic prediction is processed to redefine the pixel labels in order to obtain a binary mask with colony objects labeled as 1, which will be used to measure the size of each object and discard those not reaching or exceeding a preset minimum and maximum sizes in pixels (which would fix the thresholds to discriminate which could be a real colony and what can be noise). Refined binary masks are stored in a three-dimensional matrix, converted to boolean type and computed by means of a binary AND operation to obtain a conservative consensus given by all set of weights.

f) Resultant binary mask is then relabeled and processed by a dedicated script to identify individual colonies in close contact forming groups. To do so, each object bounding box is individually analyzed to obtain the distance profile measured from the object boundary to its innermost region. The profile is further taken as input to infer the potential number of circumferences using a local maxima peak searcher. For those objects containing more than one center, the covered pixels of each segregated colony is inferred using a watershed algorithm and their labels updated to take into account the increase of colonies in the overall counting. After that, the image is saved in a *.png file.

Once the panoptic prediction is ready, colonies are screened according to user’s criteria. To do that, detected colonies are examined individually up to three times to evaluate size, color and fluorescent intensity. Evaluation is performed by separated scripts where segments are sorted according to these properties and scored with respect to their ordinal position in the generated list. In the case of size, raw pixel area is used as evaluator to assign the related score (S_size_). In the case of color, the implemented script is designed to discriminate blue / white cells typically obtained when growing *E. coli* cells using X-Gal. To do so, a reference color (blue) is compared with the pixel-average color obtained in each segment using the HSV color model. The degree of proximity between both values is evaluated as the Euclidean distance computed from the color channel values as coordinates. We chose HSV instead RGB because under this system, colors selection is performed delimiting a cylindrical sector of the color map (recall HSV displays actual color Hue as angular coordinate, and radial and axial coordinates corresponds with saturation and value), and thus selection of a specific color and their degrees (clear or dark) is computationally performed using threshold values. On RGB, this selection involves delimiting a disjoined volume (which is less practical to formalize computationally). Colonies are then sorted according to their Euclidean distance to HSV reference color using a “dissimilarity index”: the largest the value, the more different is the evaluated color from chosen reference. The respective score (S_color_) is generated from the sorted list.

For fluorescent intensity, average intensity values are computed for each analyzed segment, and colonies are sorted in descending order to assign the fluorescence based score (S_fluor_).

After evaluating the scores of every colony in base of the required filtering criteria, a global score is computed as a weighted average of each score:

$$S_{Total}=w_{1\cdot}S_{size}+w_{2\cdot}S_{color}+w_{3\cdot}S_{fluor}$$

where w_1_, w_2_ and w_3_ are normalized weights associated to each filter criterium whose value can be chosen by the user to ponder the contribution of each criterion on the selection to perform. The selection script performs a final sorted list of colonies (in descending order) using the general mark (S_Total_). This approach allows the screening of colonies using a multicriteria selection system.

g) Positions of selected colonies are computed by transforming pixel coordinates to mm coordinates. To do that, conversion factors in x and y axis are derived by performing a calibration using a linear regression model between mm and pixel coordinates. Values are stored into a *.csv file and uploaded to OT-2.

### 2 Calibration of pixel to mm scaling function

Opentrons 2 coordinate system is given in real millimeters. Any instruction of displacement must be given in this unit. Nevertheless, positions of segmented colonies are given in pixel coordinates, because inference is based on an image. A mapping function to rescale coordinates from pixel to mm is thus necessary.

Ideally, any CCD sensor should have a constant scaling factor to link each degree of freedom (X and Y positions) in both reference systems. However, the lens of the camera may generate small distortion on the light projection from the plate to the CCD sensor, changing this relationship. We made some tests and found that a linear regression was enough to correct this bias. To calibrate the scaling function, we followed the next steps:

1- Select a plate with a low number of colonies (around 20).
2- Using a piece of paper, a pen and a ruler, find the center of the Petri Dish and mark two small lines in the upper and right borders, so you can always find a fixed orientation to look the plate. Up and right marks must be orthogonal and be aligned with the center of the plate, forming a perfect 90º angled sector (like the one draw in the piece of paper).
3- Measure the X and Y cartesian coordinates of each colony in real mm using as reference system the two orthogonal lines intersecting the centre of the plate and the orthogonal marks on the borders of the plate. You can use a rule and an erasable pen to mark reference axis in the piece of paper used in previous steps.
4- Place the Petri Dish onto the transilluminator. Using a ruler, orient the plate so the up and right marks have an orthogonal direction respect to the borders of the diffusor sides. This will ensure that the plate is oriented in the same position as before (when gathering coordinates in mm)
5- Execute COPICK functions to grab and detect colonies (steps 3 and 4 in the code commentaries).
6- Gather the region properties of variable *panoptic_completed*, which contains the labelled segments of the inference. Labels of regions (excluding the label of the agar) are stored in a vector (i.e.):

*panoptic_labeled=label(panoptic_completed)*

*info_objects_sel=regionprops(panoptic_labeled)*

*labels=np.unique(panoptic_labeled)*

*labels=labels[1:len(labels)]*

7- Then, iterate a loop using the labels to find each segment in the plate. Grab the X,Y pixel coordinates of each detected segment. Plot each segment to match each colony with stored segments.
8- Fit a linear model between pixel coordinates and mm coordinates for every dimension (X and Y axis). Apply the values to the main code.

### 3 Calibration of labware center position in pixel coordinates

As described in section 2.6 of the main manuscript, mapping the position of colonies from pixel to mm requires in advance the position of the labware center in pixel coordinates (O_lab_) to compute the pixel offset between the center of labware and the Petri Dish (ΔD_Opet-Olab_). To obtain this value, follow these instructions:

1- Take a paper sheet and place it onto the transilluminator. With a pencil or pen, mark the perimeter of the light diffusor (the flat area where Petri dish is placed) to a rectangle in vertical position. Cut the rectangle with a pair of scissors.
2- Draw the diagonals of the rectangle to find the geometrical center of the labware.
3- Place the piece of paper onto the transilluminator surface and grab a picture using the respective function
4- Open the image with an image editor and find the pixel coordinate of the intersection.
5- Update the parameters “transillum_settings['CX_labware']” and “transillum_settings['CY_labware']” in the main code.

### 4 Metrics and calculation of COPICK performance tests

Evaluation of performance tests was calculated by manual gathering of data from a dataset of 44 Petri Dishes. Related data and calculations can be found as an excel file included in the supplementary Material package. Image set containing evaluation images can be found in Github repository. Variables of interest are next detailed:

- *Plate number*: The id of the plate.
- *File name:* The name of the plate, it matches with the id number.
- *Real colony number:* Number of colonies counted manually in the plate. Groups were treated as a group of individual colonies of N elements.
- *Detectron 2 segmented colonies:* Number of segments detected by COPICK inference model
- *True positive:* Number of segments detected by COPICK and picked by the robot that were real colonies.
- *Noise / Artifacts detected & picked:* Number of segments detected by COPICK and picked by the robot that were NOT real colonies.
- *Objects_in_groups_picked_as_single (groups):* Number of segments containing groups of colonies that were detected and picked by COPICK as single colony segments.
- *Colonies_in_groups_picked_as_single (counts):* Number of individual colonies forming segments containing groups of colonies, that were picked as single colony segments. It counts the error of detecting segments composed by multiple colonies as segments with only one unique colony.
- *Objects_picked_as_group (groups):* Number segments that COPICK predicted as groups of individual colonies (they were picked more than one time), but actually they were single colonies (and should be picked once).
- *Single_colonies_picked_as_groups (counts):* Number of times that COPICK picked segments detected as group of individual colonies, when actually they were single colonies (and should be picked once). It counts the multiplicity of the error.
- *Not_processed:* Total number of colonies that were on the plate and were not processed by COPICK.
- *Not_pickable (border):* Number of colonies in close contact with the plastic border of the Petri plate that were excluded from the cropped agar region, and thus they were not present during the inference.
- *Not_detected:* Number of colonies that, although included in the image fed to the inference model, they were not detected.

Aggregated metrics contains the variables required to evaluate performance metrics. Definitions are here described:

- True positives (TP): Number of colonies detected and picked by COPICK that were actual colonies.
- False positives (FP): Counts for the sum of three source of errors, the number of segments that were noise (Noise / Artifacts detected & picked), the total number of picking errors generated in segments detected as colony groups, but actually containing a unique single colony (Single_colonies_picked_as_groups (counts)) and the number of groups picked as a unique colony that were real groups (Objects_in_groups_picked_as_single (groups)).
- False negative (FN): Number of colonies screened by the inference model, but finally not detected (not detected).
- True negatives (TN): Not apply in this case study.

Chosen performance metrics (in %) are those typically evaluated in any inference model:

- Raw Performance (screened): Number of true positive cases (TP) over total screened events.
- Raw Performance (pickable): Number of true positive cases (TP) over total detectable events.
- Accuracy: Number of correct predictions (TP+TN) over the sum of total number of positive and negative counted events (TP+TN+TN+FN).
- Misclassification rate: Complementary metric of Accuracy measuring the rate of misclassification (100-Accuracy).
- Sensitivity: Number of true positive cases (TP) over the total number of positive presented to the model (TP+FN).
- Precision: Number of true positive cases (TP) over the total number of cases classified as positive (TP+FP).
